# Alkane-priming of *Beauveria bassiana* strains to improve biocontrol of the redbanded stink bug *Piezodorus guildinii* and the bronze bug *Thaumastocoris peregrinus*

**DOI:** 10.1101/2021.07.06.451384

**Authors:** Lucía Sessa, Nicolas Pedrini, Nora Altier, Eduardo Abreo

## Abstract

Insect Epicuticle hydrocarbons (CHC) are known to be important determinants in the susceptibility degree of insects to fungal entomopathogens. Five *Beauveria bassiana* isolates were phenotypically analyzed regarding their response to CHC nutrition and their pathogenicity and virulence towards high fungal-susceptible *Thaumastocoris peregrinus* and low fungal-susceptible *Piezodorus guildinii*, which are important hemipteran pests in eucalyptus and soybean plantations, respectively. Two of these isolates, resulting the most (ILBB308) and the least (ILBB299) virulent to *P. guildinii*, were also evaluated at gene expression level after growth on n-pentadecane. *B. bassiana* most virulent isolate ILBB308 showed the lowest growth on most evaluated CHC media. However, this isolate distinctively induced most of the analyzed genes involved in CHC assimilation, cuticle degradation and stress tolerance. Virulence towards low susceptibility *P. guildinii* was enhanced in both hypervirulent ILB308 and hypovirulent ILBB299 isolates after growth on *n*-pentadecane as the sole carbon source, whereas virulence enhancement towards high susceptibility *T. peregrinus* was not observed in alkane-grown fungi. Virulence enhancement towards *P. guildinii* could be mostly explained by a priming effect produced by CHC on the induction of some genes related to hydrocarbon assimilation in ILB 205 and ILB 308, such as hydrophobin (Bbhyd2) and cytochrome P450 genes (BbCyp52g11 and BbCyp52×1), and partially by the induction of genes related to cuticle degradation (Bbchit and Bbcdep1) and stress tolerance (Bbsod1) observed only in ILB308.

## 1. Introduction

Entomopathogenic fungi, such as *Beauveria bassiana*, are used worldwide as biopesticides for pest control due to their capacity to infect arthropods by means of direct penetration through the cuticle. The overall infection process can be described in three phases. First, aerial conidia take contact with and adhere to the surface of host cuticle by non-specific hydrophobic and electrostatic interactions (Boucias et al., 1988) and by the expression of a set of hydrophobin genes (*Bbhyd1* and *Bbhyd2*) which are involved in surface hydrophobicity and adhesion, as components of the protective spore coat structure known as rodlet layer (Zhang et al., 2011). Secondly, conida germinate forming the appressorium that helps to penetrates the cuticle by a combination of mechanical pressure and enzymatic degradation (Charnley & St. Leger, 1991). The main enzymes produced are cytochrome P450s (several BbCYP families), proteases (*Bbcdep1, Bbpr1* and *Bbpr2*), and chitinases (*Bbchit1*) which help in the prosses of invasion of the hyphae to the hemocoel (Fang et al., 2005 and 2008; Zhang et al., 2009, 2010, 2011b, 2012, Pedrini et al., 2010). Finally, once in the hemocoel, conidia switch to develop as hyphal bodies evading the immune system, nurturing of hemolymph components, proliferating, disseminating and secreting toxins inside the haemocoel and various tissues, leading to the death of the insect host (Pedrini, 2018). After the insect’s death, fungal cells in the hemocoel reach the cuticle from within, escape and sporulate on the cadaver, allowing conidia to disperse and start the infection process in a new individual potentially causing epizootics.

A successful infection outcome which implies the death of the insect, depends in first instance on the penetration of the cuticle, which is composed of a thin outer layer (epicuticle) containing mainly lipids, and a thick layer (procuticle) containing chitin and proteins (Juarez et al., 1985; Blomquist et al., 1987). The epicuticular lipids are mostly hydrocarbons, i.e. *n*-alkanes, n-alkenes and methyl branched chains.

Studies have shown that *B. bassiana* can degrade a variety of hydrocarbon structures similar or identical to those of their insect host, utilizing them for energy production but also incorporating their degradation products into different fungal lipids (Napolitano & Juarez, 1997; Pedrini et al., 2010). Moreover, it is reported that nutrition on hydrocarbons enhances virulence in some *B. bassiana* strains as germination in the epicuticle is probably favored by the augmented affinity to insect-like cuticle components (Pedrini et al., 2009; Crespo et al., 2002) linked to induction of genes involved in alkane assimilation (Pedrini et al., 2010, 2013; Zhang et al., 2012). This approach seems to be getting attention in pest control management strategies with microbial control agents, and virulence gene profiling together with virulence enhancement through adaptation to growth on insect-like culture media appears to be the next step.

In Uruguay two of the most economically important insect pests are the red-banded stink bug *Piezodorus guildinii* (Westwood) (Heteroptera: Pentatomidae) and the bronze bug *Thaumastocoris peregrinus* Carpintero & Dellapé (Heteroptera: Thaumastocoridae). These pests cause significant yield losses in soybean and *Eucalyptus* plantations, respectively (Zerbino 2010; Martínez & Bianchi, 2010). Whereas chemical pesticides applied to reduce *P. guildinii* insect populations are not very successful, chemical control of *T. peregrinus* is not suitable due to the wide distribution of the insect within the canopy of the tree (Martínez & Bianchi, 2010), and also because eco-certification programs for international trade of wood products impose restrictions to the use of chemicals (https://us.fsc.org/en-us/certification). Biological control by *B. bassiana* seems to be an appropriate alternative to controlling both pests, and its effectiveness has been previously reported (Parys & Portilla, 2020; Corallo et al., 2019; Abreo et al., 2019). However, unlike for the bronze bug, no epizootics have been reported for *Piezodorus guildinii*. This suggests that virulence of pathogenic strains should be enhanced if biological control of *P. guildinii* is to be attempted. In this regard, fungal growth on an insect-like culture media, is known to augment fungal virulence. A recent study focusing on the epicuticular hydrocarbons of *Piezodorus guildinii* showed that *n*-pentadecane and *n*-tridecane are the predominant compounds of volatile short chained hydrocarbons in *P. guildinii* (Sessa et al., 2021) but not in *T. peregrinus* (Abreo et al. 2015) and could therefore be used as substrate to improve virulence of *B. bassiana* strains towards *P. guildinii*.

In this study we hypothesize that growth on hydrocarbons may serve as a virulence enhancement method for obtaining an effective *B. bassiana*-based biopesticide against *P. guildinii*. The aims of the present work were to evaluate i) the effect of hydrocarbon nutrition on vegetative growth strains and virulence of five *B. bassiana* towards *P. guildinii* and T. peregrinus, and ii) the effect of *n*-pentadecane -used for virulence enhancement- on gene expression regarding relevant to infection genes of two contrasting *B. bassiana* strains.

## 2. Materials and Methods

### 2.1 Entomopathogen fungal strains

Five *Beauveria bassiana* isolates were evaluated in this study. Isolates ILB308, ILB299 and ILB205 were retrieved from INIA-Las Brujas Fungal Collection (ILB), Fi1728 was provided by Laboratorio de Micología-Universidad de la República, and strain GHA is a commercial strain (Laverlam International, Butte, MT). Isolates were revitalized on mealworms (*Tenebrio molitor*) and stored on PDA tubes and plates at 4°C.

### 2.2 Fungal cultures and spore suspension preparation

Every spore suspension used in this study was obtained from aerial spores harvested and resuspended with Twee*n*-saline solution (0.1% Tween 80 and 0.85%NaCl), filtered using autoclaved gauze and adjusted to its final concentration using a Neubauer chamber. Spore viability (%) of each isolate suspension was assessed by plating an aliquot of 150 μl of a 1/10 diluted suspension (2 × 10^6^ spores/ml) on a water agar plate (WA) and incubated for 24 hours at 26°C. After the incubation period, viability was evaluated as the number of viable spores out of the total number of counted spores, multiplied by 100. Viability was measured in three areas per plate, with one hundred total spores counted each time. Spores were produced either on potato dextrose agar (PDA), complete media (CM) or on minimal media (MM) supplemented with HC. CM plates contained 0.4 g KH_2_PO_4_, 1.4 g Na_2_HPO_4_, 0.6 g MgSO_4_.7H_2_O, 1.0 g KCl, 0.7 g NH4NO_3_.7H_2_O, 10 g glucose, 5 g yeast extract and 15g agarose in 1000 ml of distilled water and MM contained the same as CM without the glucose and yeast extract, and supplemented with *n*-alkanes (Thermo Fisher Scientific, USA) as carbon source (Pedrini et al. 2010).

### 2.3 Rearing of insect colonies

Individuals of *T. peregrinus* were obtained from an indoor mass rearing maintained on fresh shoots of *Eucalyptus tereticornis* Sm. in Erlenmeyer flasks (Martínez et al., 2014). Nymphs were placed on aluminum framework cages covered with a mesh screen and were checked weekly for adult emergence. Adult individuals were sexed and relocated into similar cages prior to the inoculation.

Individuals of *P. guildinii* were obtained from an indoor mass rearing maintained at 24 ± 1°C on fresh soybeans, green beans, and water. Nymphs were placed on transparent plastic boxes and checked daily until adult emergence. Adults were sexed and placed on plastic cages prior to the bioassay.

### 2.4 Pathogenesis and virulence assay towards *P. guildinii* and *T. peregrinus*

A preliminary pathogenicity assay was conducted using PDA-grown fungal conidia. Isolates ILB308, ILB299, ILB205, Fi1728 and GHA were grown on PDA plates and conidia were harvested after 14 days of incubation at 26°C. Suspension concentration was adjusted to 2 × 10^7^ viable spores/ml.

Twenty laboratory-reared adults of *Piezodorus guildinii* insects (10 males and 10 females) were inoculated with each spore suspension. Insects were immersed in the *B. bassiana* spore suspension for 5 seconds and immediately placed individually on a 55mm Petri dish containing sterile paper, water, and soybean grains for insect nutrition during incubation. Controls were immersed in water/Twee*n*-saline solution. In total 120 insects were used. All treated insects were incubated at 24 ± 1°C for 12 days and insect mortality was assessed and recorded daily. Each insect cadaver was placed on a humid chamber at 25°C for fungal development. This entire trial was repeated three times, evaluating a total of 360 *Piezodorus* insects.

Fifty laboratory-reared adults of *T. peregrinus* (25 males and 25 females) were placed on bouquets of fresh leaves of Eucalyptus tereticornis Sm. and kept in voile mesh cages, according to Martínez et al., 2014. Each conidial suspension was applied by spraying onto the bouquets to the point of run-off. Three bouquets were sprayed per fungal strain. Control bouquets were sprayed with water/tween-saline solution. Cages were kept for 10 days under controlled conditions of temperature and moisture (23 ± 2°C and 80% relative humidity) under 12-h light/dark period. Mortality of insects was assessed and recorded daily. Each insect cadaver was superficially disinfected in alcohol 70% for 5 s, washed twice with sterile distilled water, dried on sterile filter paper, placed in individual humid chambers, and incubated at 26 °C for fungal development. This entire trial was repeated once, evaluating a total of 18 cages containing 900 insects.

Cumulative survivalship curves were constructed for both bioassays. The median lethal time (MLT) was the parameter used for assessing virulence in *T. peregrinus* as all treated insects reached 100% mortality. MLT was estimated as Σ (days_n_ × dead insects_n_)/total dead insects for (Moore et al., 1995). Abbot’s corrected mortality (Abbot, 1925) was the parameter used for assessing virulence in *P. guildinii* inoculated insects because none of the treatments reached total mortality.

### 2.5 *Beauveria bassiana* growth on *n*-alkanes

The preference of the *B. bassiana* isolates for *n*-alkanes was evaluated by assessing vegetative growth on MM culture media supplemented with different *n*-alkanes and CM. Four *n*-alkane substrates were evaluated, selected according to their relative abundance on the epicuticular layer of each of the two insects, *n*-tridecane, *n*-pentadecane and two blends. The long chain hydrocarbon mix (LC-HC) resembled the relative abundance of epicuticular hydrocarbons found in the soybean stink bug and consisted of a mixture of *n*-tridecane, *n*-pentadecane and *n*-octacosane in proportion 1:0.9:0.1. The very long chain hydrocarbon mix (VLC-HC) resembled the relative abundance of epicuticular hydrocarbons found in the bronze bug and consisted of a mixture of *n*-tetracosane and *n*-octacosane in equal proportions (1:1). To assess vegetative growth, a 20 μl droplet of 6 × 10^7^ viable spores/ml of each suspension was inoculated in the center of a petri dish with CM or MM solid media. Each strain and condition were repeated three times. Plates were incubated at 26°C and diameter growth was recorded on the sixth day post inoculation.

### 2.6 Expression of genes involved in adhesion, hydrocarbon assimilation, pathogenesis and oxidative stress tolerance using qRT-PCR

Levels of differential expression of 10 relevant to infection genes (Table1) in CM and in MM supplemented with *n*-pentadecane were investigated, to assess the gene expression patterns during growth on hydrocarbon. Isolates were grown on solid MM supplemented with *n*-pentadecane (C15, 1 % final concentration) and CM plates, covered with a cellophane membrane to avoid alkane interference with the RNA integrity and qRT-PCR protocol. Plates were incubated for 9 days at 26°C. Total RNA was extracted on day 3, 6 and 9 employing the RNeasy plant mini kit (Qiagen). RNA was quantified using Nanodrop (Thermo Fisher) and integrity and purity were evaluated on an agarose gel 1% electrophoresis. Samples were treated with Turbo DNase (Invitrogen, Thermo Fisher Scientific) and EDTA. Purified RNA was transcribed into cDNA using iScript cDNA Synthesis Kit (Bio-Rad), and stored at −20 °C until qRT-PCR analysis. The assay was performed three times (three biological replicates).

qRT-PCR analyses were performed using iQ SYBR Green Supermix (Bio-rad). PCRs were performed using Quant Studio 3 (Applied Biosystems, Thermo Fisher), using the following conditions: denaturation at 95 °C for 3 min, followed by 40 cycles of 95 °C for 15 seconds and 60 °C for 30 seconds. Data was analyzed using the Quant Studio design and analysis software version 1.4.3. Transcripts were amplified using 10 gene-specific primers detailed in table 1. For normalization the glyceraldehyde-3-phosphate dehydrogenase (GAPDH) gene from *B. bassiana* was used (Table 1). The qRT-PCR analyses were performed in triplicate for each of the three independent biological replicates.

**Table 1.**
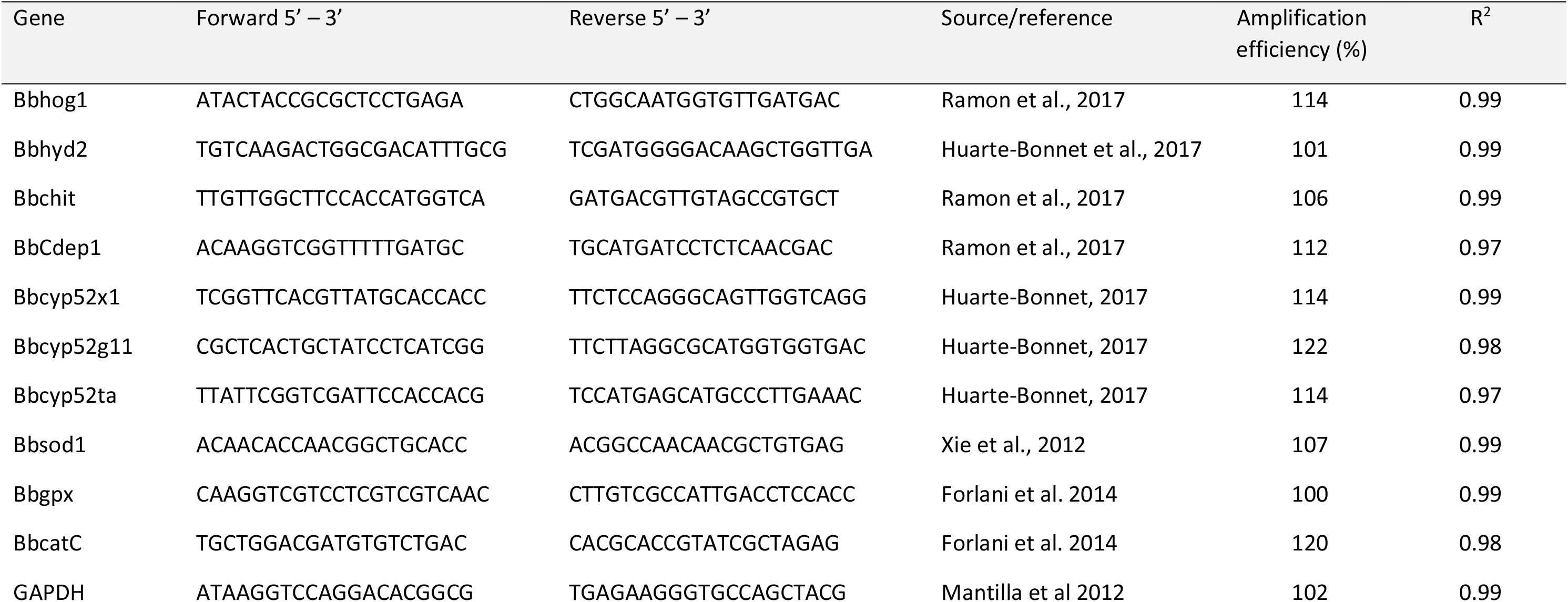
PCR amplification primer sequence and amplification efficiency

To assess the primer’s amplification efficiency, standard curves were constructed by plotting the log of the cDNA dilution values versus CT values. Serial 5-fold dilutions of cDNA samples were used as templates for each qRT-PCR reaction. Each reaction efficiency was calculated as E = 10 ^(−1/slope)^, using the slope of each curve (table 1). The relative expression levels of each gene were calculated using the 2^−ΔΔCt^ method (Livak).

### 2.7 Virulence enhancement of *B. bassiana* strains grown on *n*-alkane towards *P. guildinii*

and *T. peregrinus*

The virulence enhancement bioassays were conducted using 2 × 10^7^ viable spores/ml solutions obtained from aerial conidia of ILB308, ILB299, ILB205, Fi1728 and GHA isolates grown on CM and MM amended with *n*-pentadecane plates. Because ILB299 showed reduced sporulation, it was evaluated at a lower concentration once on *T. peregrinus* using 2 × 10^6^ viable spores/ml. Inoculation and evaluation procedures, including humid chambers, were as described on the previous bioassays, and each assay was repeated twice using 10 individuals per condition. Cumulative survivalship curves were constructed for both bioassays, using the same parameters as described in section 2.4.

## Results

### *B. bassiana* virulence towards *T. peregrinus* and *P. guildinii*

Pathogenicity baseline of the five fungal strains was determined to select a high and a low virulent strain (hyper and hypo virulent) for future evaluations and to define the time frame for the gene expression experiments.

*Thaumastocoris peregrinus* proved to be very susceptible to all *B. bassiana* strains, as survivorship in all treated insects reached 0% by the end of the evaluated period (Fig. 1a). Mean lethal time was estimated among pathogenic strains. ILB205, GHA and ILB308 were the most virulent with MLT values of 5.21±0.05, 5.84±0.06 and 5.89±0.12 days, respectively (Supplementary T1).

**Figure 1.**
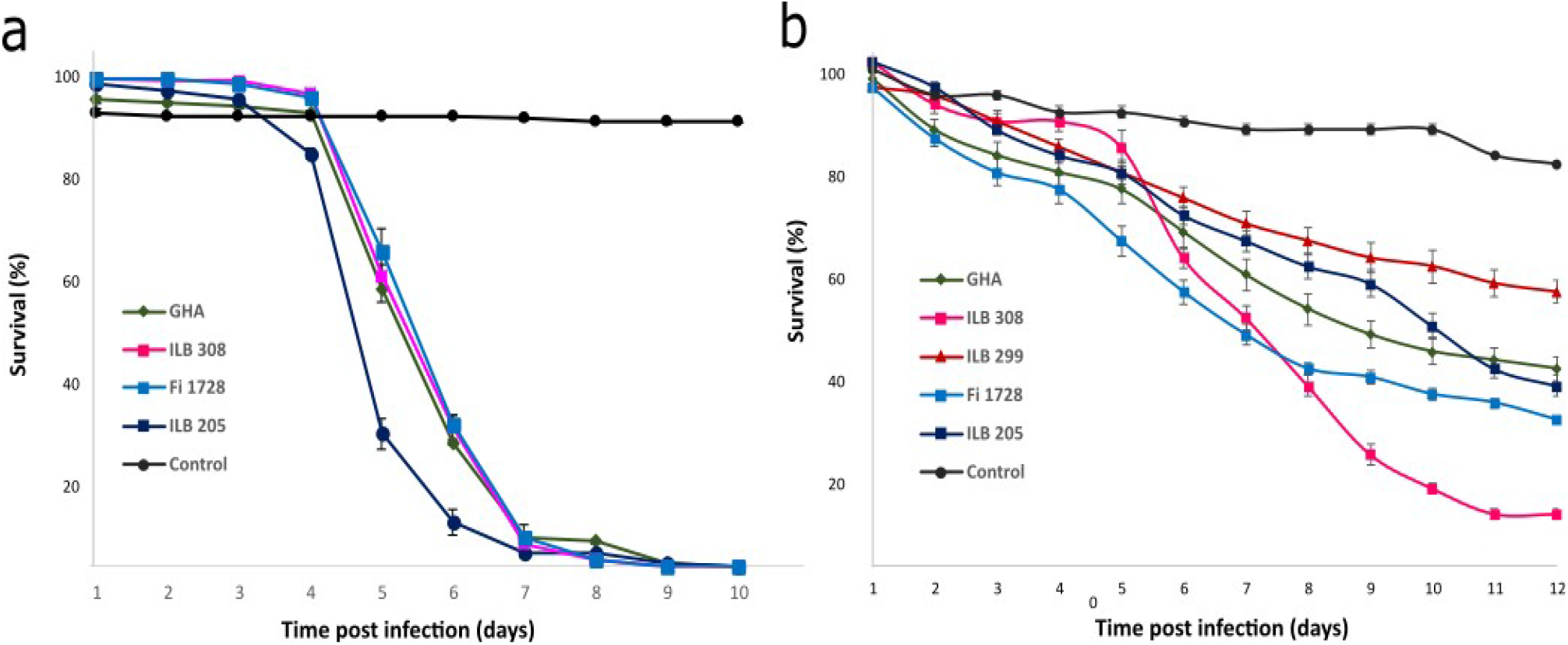
Cumulative daily survival of (a) *Thaumastocoris peregrinus* and (b) *Piezodorus guildinii* exposed to different PDA grown *Beauveria bassiana* isolates. Data shows means ± SD form corresponding replicates.

*Piezodorus guildinii* insects showed less susceptibility towards *B. bassiana* infection, although isolates proved to be pathogenic as they differed from the mortality rates observed in the control treatment (Figure 1b). No strain provoked total mortality by the end of the assay. Abbots corrected mortality (ACM) on the assay’s end day was calculated to compare virulence among strains. ACM values showed that strains ILB308 and ILB299 had the highest and lowest mortality rates, 0.91 ± 0.09 and 0.26 ± 0.19, respectively (Supplementary T 2).

### *Beauveria bassiana* growth on *n*-alkanes

All strains evaluated were able to utilize synthetic *n*-alkanes as the sole carbon source, as evidenced on the vegetative growth evaluation. As expected, the preferred culture media was CM as all strains showed higher total growth (Table 2), as well as more abundant aerial mycelia (Fig. 2). Overall, the strains with higher vegetative growth on any HC or their combinations were Fi1728, IL205 and ILB299, and strain GHA together with ILB308 had the lowest (Fig. 2, Table 2). Morphologically strain ILB308 differed from the rest of the *B. bassiana* strains, showing scarce aerial mycelial development (Fig. 2).

**Table 2.** Colony diameter (cm) of *B. bassiana* strains measured after 6 days of growth on complete media (CM) and minimal media supplemented with different *n*-alkanes

|  | HC-supplemented MM |  |  |  | CM |
| --- | --- | --- | --- | --- | --- |
|  | *n*-tridecane | *n*-pentadecane | LC-HC | VLC-HC |  |
| Fi1728 | 2.33 ± 0.23 c AB | 2.68 ± 0.08 bc A | 2.76 ± 0.12 b A | 2.56 ± 0.12 bc A | 3.36 ± 0.03 a BC |
| ILB299 | 2.53 ± 0.09 b A | 2.45 ± 0.27 b AB | 2.56 ± 0.03 b A | 2.53 ± 0.03 b A | 2.98 ± 0.03 a D |
| ILB308 | 2.13 ± 0.11 b BC | 1.91 ± 0.24 b B | 1.90 ± 0.43 b A | 1.98 ± 0.03 b B | 3.53 ± 0.03 a B |
| ILB205 | 2.53 ± 0.04 b A | 2.40 ± 0.35 b AB | 2.44 ± 0.36 b A | 2.42 ± 0.15 b A | 3.16 ± 0.06 a CD |
| GHA | 1.98 ± 0.06 b C | 2.00 ± 0.21 b B | 2.00 ± 0.51 b A | 1.99 ± 0.17 b B | 3.97 ± 0.17 a A |

Means (± SD) followed by different letters denote statistical differences (Tukey's HSD test, P<0.05) Capital letters denote differences in columns, lower case letters indicate differences in rows

**Figure 2.**
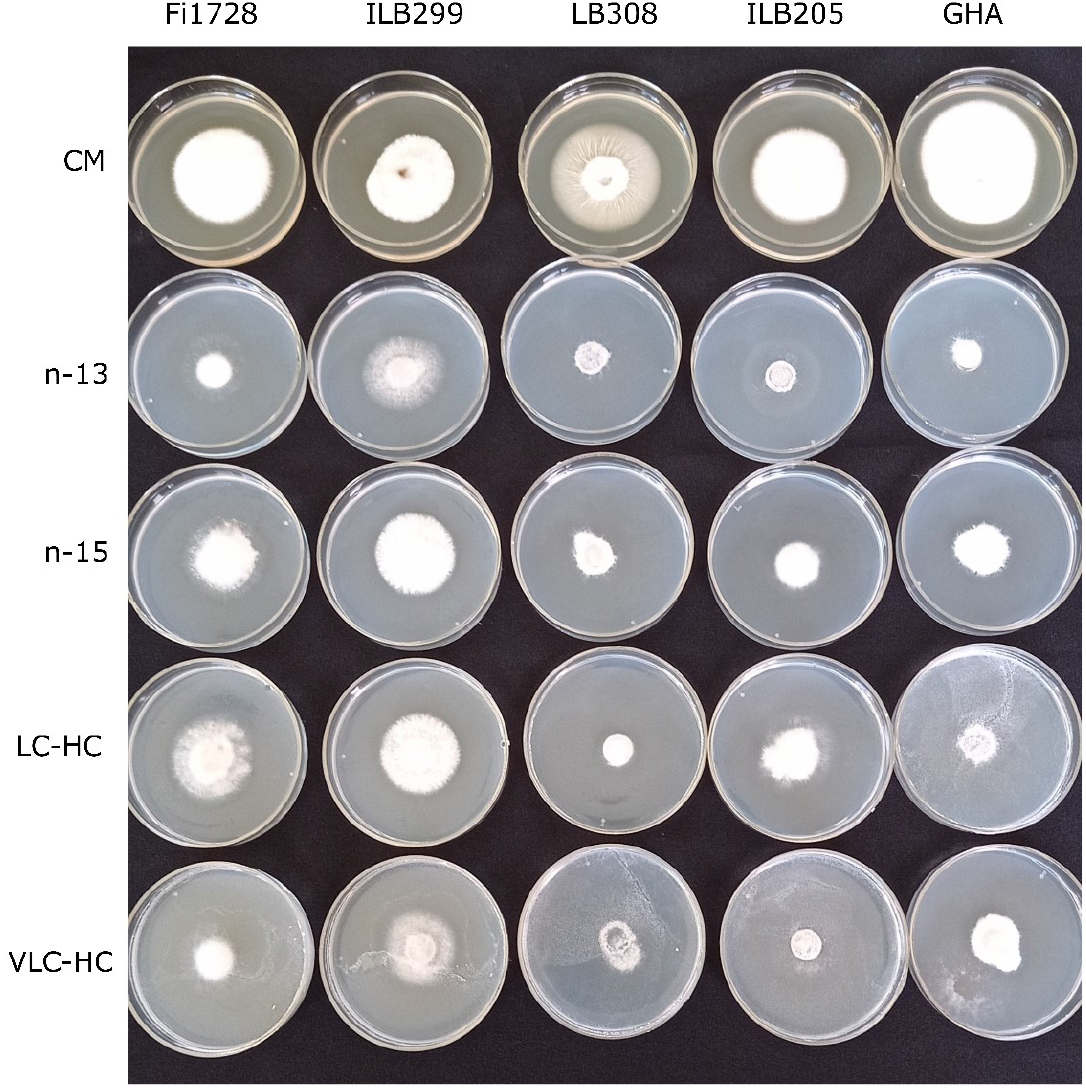
Vegetative growth evaluation after 6 d of incubation at 26±2 °C. A 20μl droplet of *Beauveria bassiana* conidial suspensions were inoculated on complete medium and minimal medium supplemented with different *n*-alkanes.

### Expression of *B. bassiana* genes involved in hydrocarbon assimilation (*Cyp52g11, Cyp52t1* and *Cyp52×1*)

Regarding genes involved in hydrocarbon assimilation, a delay in the induction was observed in the low virulence strain ILB299. This strain showed under expression of the three genes on the first day of the evaluation. Later, on the 6^th^ day *Bbcyp52g11* and *Bbcyp52×1* genes exhibited very low overexpression of 1.96±0.46 and 1.96±0.66 foldchange, respectively, while *Bbcyp52ta* remained under-expressed (Fig. 3). On the 9^th^ day *Bbcyp52g11* and *Bbcyp52ta* reached maximum overexpression of 2.11±0.82 and 1.61±0.83 foldchange, respectively, while *Bbcyp52×1* decreased to 0.79±0.27 (Fig. 3). At the beginning of the evaluation hyper virulent strain ILB308 exhibited overexpression of the *Bbcyp52×1* gene involved in hydrocarbon assimilation in 1.29±0.60 foldchange (Fig. 3). Later, expression of this gene, together with *Bbcyp52×1* reached a maximum, with foldchange values of 7.27±2.70 and 3.20±0.69, respectively. *Bbcyp52ta* remained over expressed.

**Figure 3.**
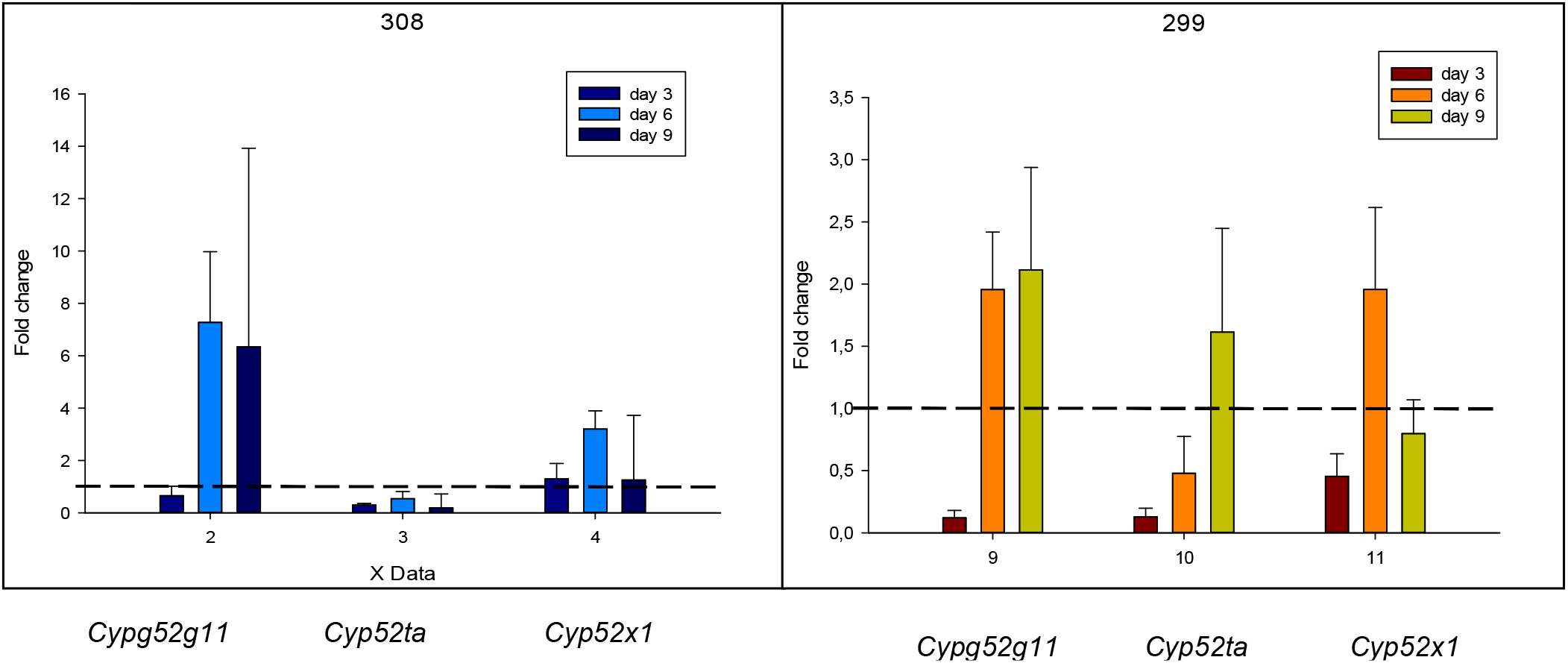
Expression pattern of hydrocarbon degradation and assimilation genes in *Beauveria bassiana* strains ILB308 and ILB299 grown on *n*-pentadecane. The dashed line shows an expression ratio = 1, indicating no expression change due to *n*-pentadecane nutrition. Values are means ± SE of three biological replicates.

### Expression of *B. bassiana* genes involved in host adhesion and appressorium formation (*Bbhyd2* and *Bbhog1*) and cuticle degradation (*Bbchit, Bbcdep1*)

The four evaluated genes (*Bbhyd2*, *Bbhog1*, *Bbchit* and *Bbcdep1*) were found under-expressed in the hypo-virulent strain ILB299 on day 3 of evaluation (Fig. 4). In fact, cuticle degradation genes remained under-expressed during the entire evaluation. The highest over-expression was observed in the host adhesion gene *Bbhyd2* on day 6 which remained so until day 9 (5.83±2.61 and 3.15±1.02 on day 6 and day 9, respectively). The appressorium formation gene (*Bbhog1*) had 1-foldchange expression on day 6 and 9 of the evaluation, meaning that *n*-pentadecane culture medium had no effect on the expression of these gene in ILB299 strain. In contrast, the hypervirulent strain ILB308 had not only the adhesion gene (*Bbhyd2*), but also the cuticle degradation gene Bbdcep1 overexpressed since day 3 (Fig. 4). On day 6 all the genes evaluated, except for Bbdcep1, including the appressorium formation gene (*Bbhog1*) and the chitinase coding gene (*Bbchit*) were overexpressed, and some remained overexpressed up until the final day of the evaluation.

**Figure 4.**
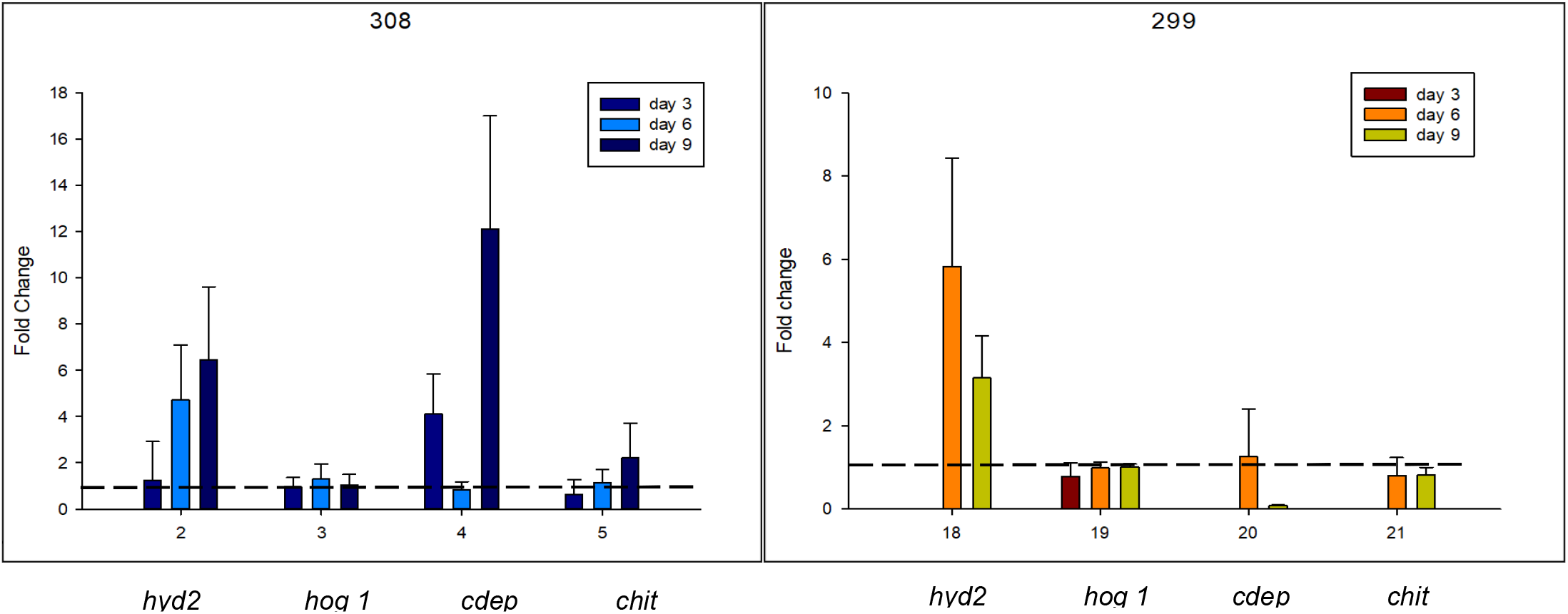
Expression pattern of host adhesion, appressorium formation and cuticle degradation genes in *Beauveria bassiana* strains ILB308 and ILB299 grown on *n*-pentadecane. The dashed line shows an expression ratio = 1, indicating no expression change due to *n*-pentadecane nutrition. Values are means ± SE of three biological replicates.

### Expression of *B. bassiana* genes involved in the antioxidant defense system (*BbcatC, Bbsod1* and *Bbgpx*)

Superoxide dismutase gene (*Bbsod1*) was slightly overexpressed during the entire evaluation period in both strains (Fig. 5). Also, in both strains *BbcatC* gene expression showed a peak on day 6 and on day 9 of the evaluation. The glutathione peroxidase gen (*Bbggpx*) was under-expressed during the entire evaluation period in the low virulence strain ILB299 and clearly over-induced on strain ILB308 up from day 6 forward, with foldchange values of 2.92±1.47 and 2.78±1.30 on the 6^th^ and 9^th^ day of the evaluation, respectively.

**Figure 5.**
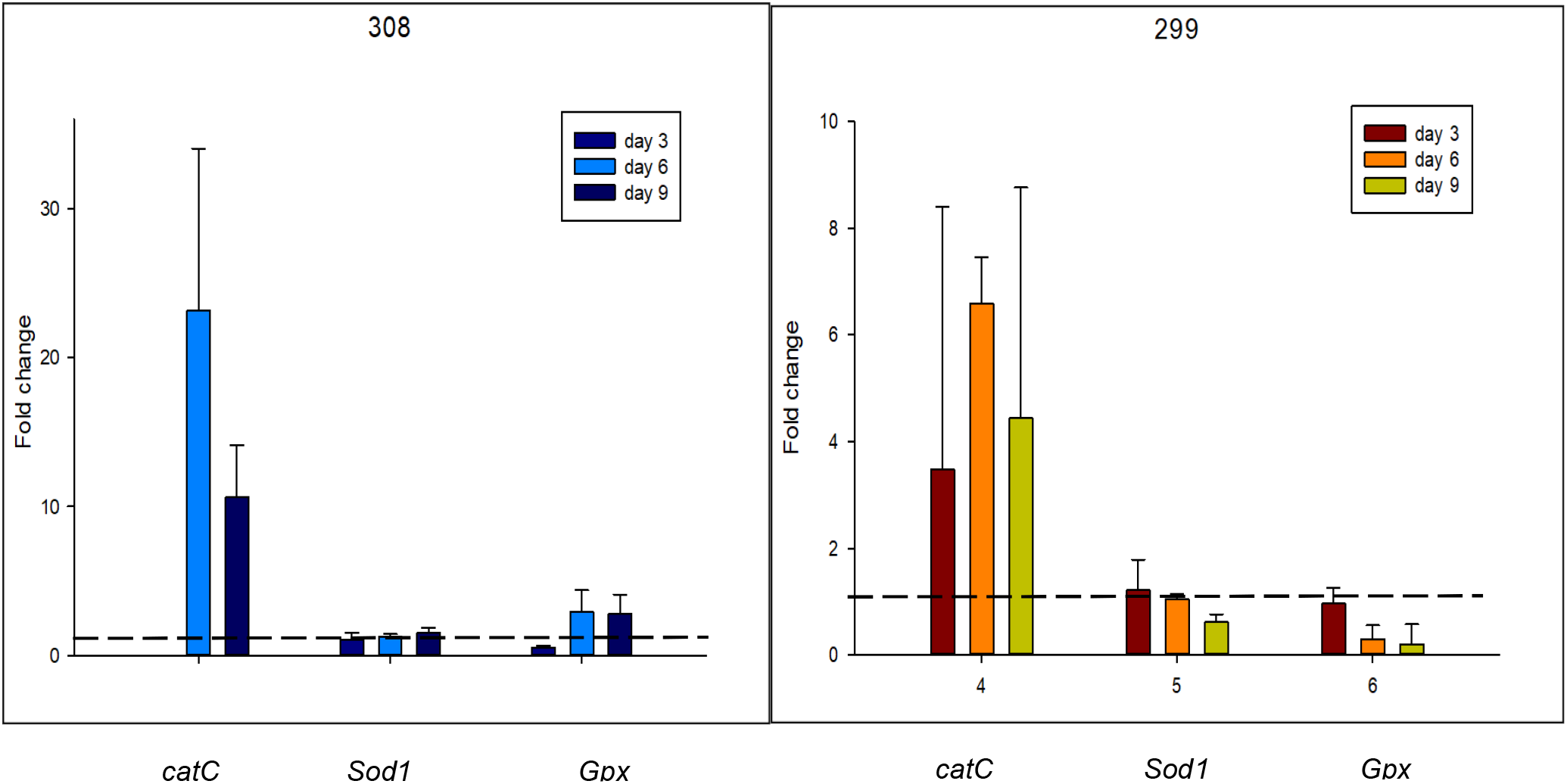
Expression pattern of the antioxidant defense system genes in *Beauveria bassiana* strains ILB308 and ILB299 grown on *n*-pentadecane. The dashed line shows an expression ratio = 1, indicating no expression change due to *n*-pentadecane nutrition. Values are means ± SE of three biological replicates.

### Virulence enhancement evaluation of HC grown *B. bassiana* strains towards *T. peregrinus* and *P. guildinii*

On *T. peregrinus*, all *B. bassiana* strains proved to be highly virulent, as insects showed very low survival percentages by the end of the assay compared with the mock control (Supplementary F1). There was no virulence enhancement caused by growth of the isolates on *n*-alkanes as no significant differences were observed between the estimated MLT of each strain grown on *n*-pentadecane compared with the same strain grown on CM medium (Supplementary T3). Figure 6 shows both hyper- and hypo-virulent strains (ILB308 and ILB299, respectively) survivalship curves, the latter curve being less pronounced due to a lower concentration of the conidial suspension inoculated.

**Figure 6.**
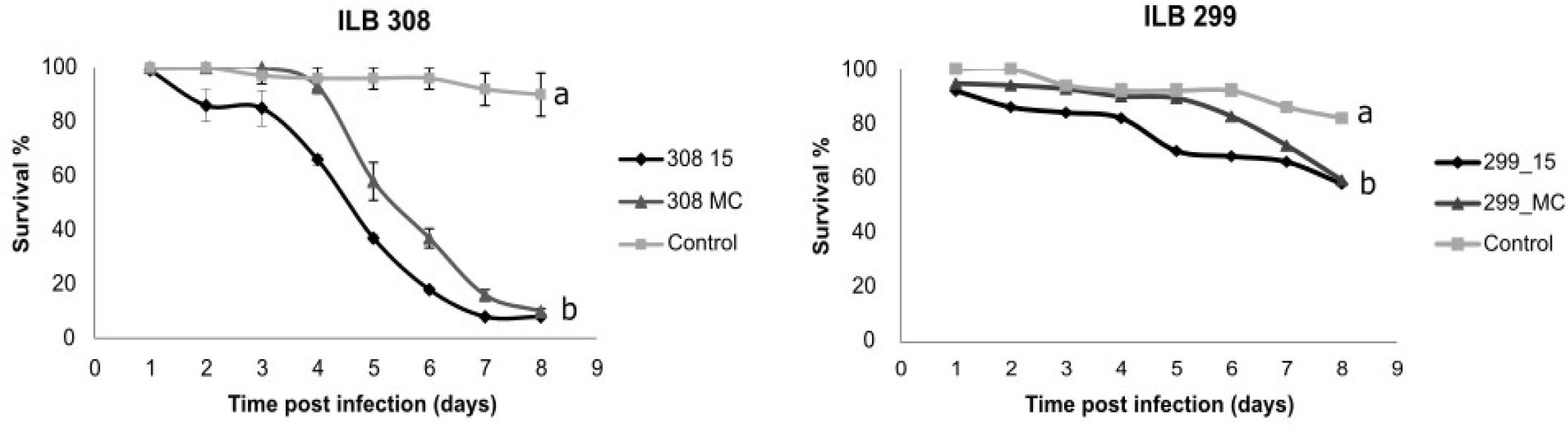
Survivalship of *T. peregrinus* infected with *B. bassiana* grown on *n*-pentadecane (15) and MC. a conidial suspension of hypervirulent strain ILB308 2×10^7^ conidia/ml. b conidial suspension of hypo-virulent strain ILB299 2×10^6^ conidia/ml.

On *Piezodorus guildinii* inoculated insects, the *n*-alkane nutrition effect on virulence enhancement was evidenced. Strains ILB308, Fi1728 and ILB299 showed an augmentation on virulence, being significantly higher in strains grown on *n*-pentadecane than CM medium (Fig. 7, Supplementary F2). Strains GHA and ILB205 showed no significant differences in mortality between conidia grown on *n*-pentadecane and conidia grown on CM medium (Supplementary T4).

**Figure 7.**
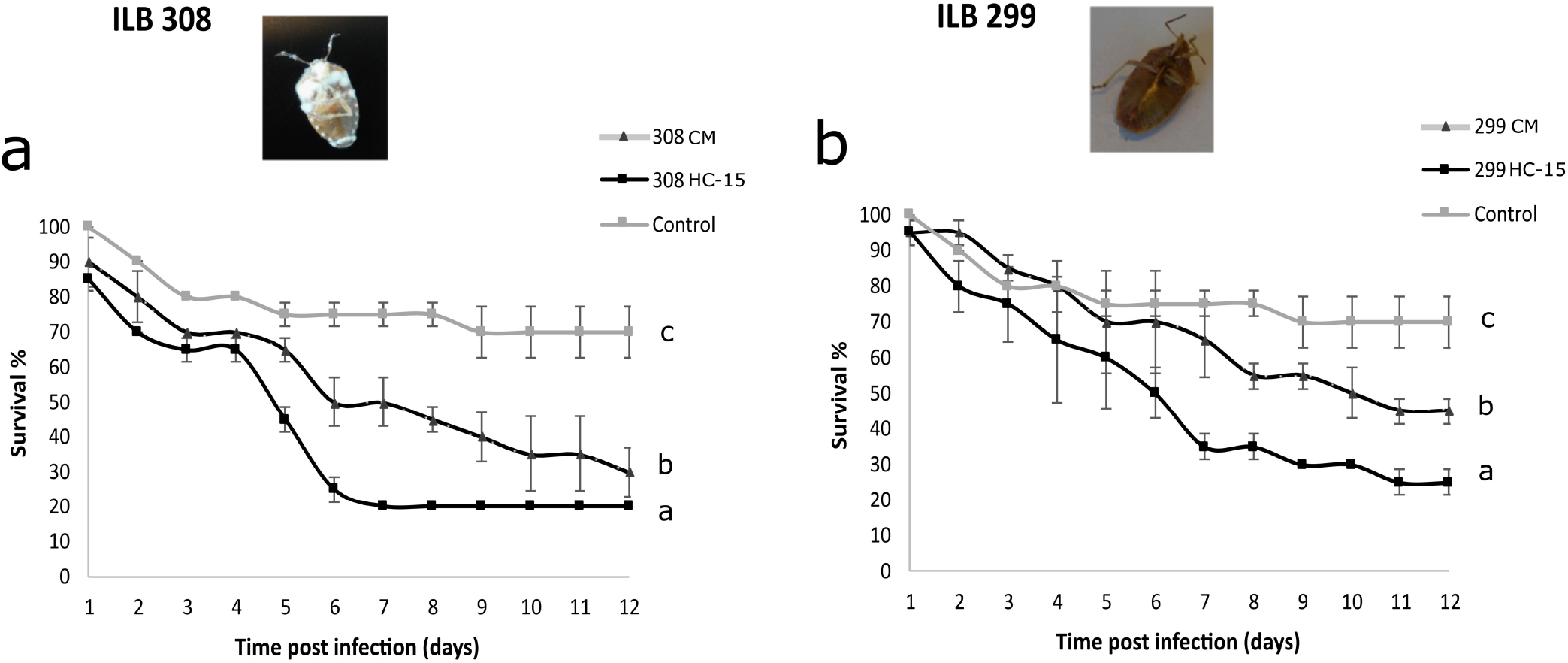
Survivalship of *P. guildinii* infected with *B. bassiana* spore suspension of 2×10^7^grown on *n*-pentadecane (HC-15) and CM. Different letters designate significant differences between survival curves. Insets: Cadavers of *P. guildinii* after 10 days post-inoculation with *Beauveria bassiana* isolates

Sporulation on dead cadavers of *T. peregrinus* in the humid chambers was evident in almost all treated insects inoculated with ILB308, ILB205 and GHA grown on both culture media. Cadavers on humid chambers of ILB299 and ILB1728 inoculated insects showed aerial mycelial growth rather than conidia.

On *P. guildinii* dead insects, sporulation in cadavers was rather scarce. Strain ILB308 showed the highest levels of mummified cadavers covered in *B. bassiana* conidia while ILB299 mycelium was barely observed on dead insects (figure 7, insets).

## Discussion

The interaction of the entomopathogen *B. bassiana* with insect pests leads to insect death when the fungal spore is firstly able to adhere and penetrate the insect cuticle and then progress to the haemocoel, where it must avoid host response, multiply and invade the insect cavity, ultimately killing the host. *Piezodorus guildinii* and *Thaumastocoris peregrinus*, two hemipteran insect pests with different epicuticle composition and susceptibility towards *B. bassiana*, were studied in their interaction with five *B. bassiana* isolates showing a wide range of virulence. We focused on the response of a hyper virulent and a hypo virulent *B. bassiana* isolate to hydrocarbons that are an integral part of the insect’s epicuticle and a first defense barrier against fungal infection. The ultimate objective was to explore the possibility of virulence enhancement through manipulation of the interaction between the fungus and the insect epicuticle components, in order to obtain a biocontrol agent capable of infecting both insect pests. This is important for the soybean bug *Piezodorus guildinii*, which is known for its low susceptibility towards fungal infection under field conditions.

Both hemipterans showed different susceptibilities towards the *B. bassiana* isolates used in this study. Inoculated *Thaumastocoris peregrinus* insects reached 100% total mortality with all evaluated *B. bassiana* isolates. This was previously noted by Corallo et al. (2019) when comparing to other entomopathogen species, such as Isaria farinosa, Lecanicillium lecanii and Paecilomyces variotii, *B. bassiana* showed the greatest virulence, killing more individuals in shorter time. On the contrary, the red-banded stink bug *P. guildinii* showed to be more recalcitrant to fungal infection, with different isolates showing different mortality percentages. The highest mortality percentage recorded was 90%, reached by the hyper-virulent strain ILB308, and the lowest was 46.7% shown by the hypo-virulent strain ILB299. Lower mortality was obtained by Parys & Portilla (2020) where at day 10 since inoculation, *P. guildinii* insects inoculated with two *B. bassiana* isolates reached 70 % mortality.

The hypothesis tested in this study was that the capacity to grow on CHC was positively co-related with virulence, since epicuticular CHC are the first barrier a spore must overcome in order to invade the insect host. Because growth on hydrocarbons is known to impose cellular stress (Cray et al., 2015, Huarte-Bonnet et al., 2014), the lower vegetative growth as well as less abundant mycelia observed on all isolates grown on MM supplemented with *n*-alkanes compared with CM medium was expected (Crespo et el., 2000). However, when comparative growth on different CHC was evaluated, ILB308-regarded as the most virulent isolate against the most resistant insect *P. guildinii*-showed the lowest vegetative growth together with the commercial isolate GHA. Isolates Fi1728 and ILB205, of intermediate virulence towards *P. guildinii*, showed higher radial growths, but in general lower than the least virulent isolate ILB299. Morphologically, ILB299 and FI1728 produced more abundant aerial mycelia on all evaluated culture media. In conclusion, no positive correlation between vegetative growth on CHC and virulence was found.

Interestingly, when analyzing the gene expression of the hypervirulent isolate ILB308 and hypo virulent isolate ILB299 grown in *n*-alkane, in relation with adherence hydrocarbon assimilation, cuticle degradation and antioxidant defense, it was clearly seen that in general, in the hypervirulent isolate the genes involved in those functions were up-regulated earlier and in a more pronounced manner than the hypo virulent strain. These results are in concordance with previous reports where early infection reported genes are up regulated in most virulent strains (Zhang et al., 2011, Wang et al., 2017). The apparent contradiction between colony growth in CHC, which was lower in ILB308, and the expression of genes related with CHC assimilation, which was higher, could be explained by the expression of other genes that could be more directly related to growth or sporulation that were not evaluated in this experiment (Zhang et al., 2010; Dias et al., 2008; Khoury et al., 2019).

Regarding the effect of *n*-alkane nutrition on the expression of virulence related genes, the most virulent isolate overexpressed genes related to virulence and stress defense earlier and in a more sustained fashion than the hypo virulent strain when grown on *n*-pentadecane. Isolate ILB308 overexpressed the *Bbhyd2* gene -coding for hydrophobi*n*- at day 3, while it was silent in the hypovirulent isolate ILB299 on the same day. Hydrophobins are necessary for the fungus to adapt to the hydrophobic wax-rich epicuticular surface and their early overexpression in ILB308 might imply better chances for the progression of the pathogenesis for ILB308 than hypovirulent ILB299. This result is in concordance with Huarte et al. (2017) where high induction of the genes coding for hydrophobins were found in hydrocarbo*n*-grown conidia.

The single-copy genes Bbcdep1 (Fang et al., 2005; Romon et al., 2017), coding for a subtilisi*n*-like protease, and *Bbchit*1 coding for a chitinase were overexpressed in strain ILB308 at day 3 and 9 of the evaluation, respectively. These two genes were not induced in isolate ILB299. As single copy genes, it seems reasonable that their relative expression levels were directly correlated with virulence. Therefore, when ILB308 grew on *n*-alkane, a priming of virulence genes related with degradation of proteins and chitin was observed, which could explain the baseline virulence of this isolate towards *P. guildinii* and partially its increased virulence after growth on pentadecane. Interestingly, a fusion protein with protease and chitinase activity based on these two genes was designed to enhance the virulence of *B. bassiana* (Fang et al. 2009). With proteins and chitin being the most abundant/important components of the cuticle, the overexpression of these two genes in ILB308 grown on *n*-alkane could be mimicking the effect obtained by the transformed strain in that study.

Regarding Bbcdep1 overexpressed in ILB308 and silent in ILB299, a recent study by Gao et al. (2020) found CDEP1 to be correlated to higher fungal outgrowths on insect cadavers and their conidial yield, implicating a link of this protease to fungal survival, dispersal, and infection cycle in host habitats. So, the differences observed in our study in the expression of this gene, particularly the under-expression observed in ILB299 may be correlated to the incapacity of this isolate to breach the insect cuticle from the inside and sporulate on dead cadavers.

Because growth on alkanes causes major changes in fungal metabolism (Crespo et al. 2000) and a scenario of oxidative stress is caused by the accumulation of reactive oxygen species, the induction of antioxidant genes and enzymes to overcome this situation can be understood as another example of fungal higher fitness (Huarte-Bonnet et al. 2015). In this study both isolates showed a clear induction of the stress tolerance gen BbcatC and a slightly induction of *Bbsod1* gen. The GSH system gene Bbgpx (selenium-glutathione peroxidase enzyme) was differentially expressed, being induced in the hypervirulent isolate ILB308 and under induced in hypovirulent isolate ILB299. In this sense, because gpx expression is regulated to maintain catalase activity (Baud et al., 2004), the induction of this gene may offer an advantage in the oxidative response of the hypervirulent isolate ILB308.

Enzymes involved in terminal hydroxylation of *n*-alkanes and fatty acids (P450alk) are expressed from genes belonging to the CYP52 family (Tanaka et al., 1982; Pedrini et al., 2013), which act on a wide variety of endogenous and xenobiotic molecules, including alkanes, a major constituent of many insect cuticles. Here we evaluated the expression of Cyp52g11, Cyp52ta and Cyp52×1 genes involved in hydrocarbon assimilation. Results show that *Bbcyp52g11* and BbCyp52×1 genes were induced in both fungal isolates, although induction levels were higher and earlier in time on isolate ILB308.

It is remarkable that the hyper and hypo virulent strains, showing the previously described differences in original virulence, growth on CHC and gene expression, were similarly affected in their capacity to infect and kill *P. guildinii* insects after growth on *n*-pentadecane. The HC15 priming effect on virulence in both strains seems linked to the overexpression of the genes belonging to the CYP52 family evaluated on this work Bbcypg11 and Bbcypx1, and relatively independent from the observed induction of *Bbchit*, Bbcdep1 and Bbgpx genes, which were clearly overexpressed in the hypervirulent ILB308 and not in the hypo virulent strain ILB299. This finding is in concordance with reports that suggest that the higher virulence observed in hydrocarbon grown strains is due to induction of alkane assimilation genes (Pedrini et al., 2010, 2013; Zhang et al., 2012). In other words, the genes whose overexpression could explain the difference in original virulence between ILBB308 and ILBB299 are not the ones that can be signaled as the responsible for the virulence enhancement observed in both strains after growth on *n*-pentadecane. Rather, the overexpression of genes related to CHC assimilation, observed in ILB308 and ILB299 when grown on *n*-pentadecane, could be responsible of the virulence enhancement observed in both isolates.

In the case of *T. peregrinus*, a rather susceptible insect pest, conidia harvested from the hyper and hypo virulent strains grown on *n*-pentadecane, had the same virulence as conidia harvested from CM. We hypothesize that due to the high susceptibility of this hemipteran towards the tested fungal isolates, the priming effect could be limited as there was not room for improvement. Yet, another hypothesis could be that the HC selected for priming was a *P. guildinii* specific alkane, and possibly the effect was somewhat directed towards this hemipteran.

We have shown that hypervirulent isolate ILB308 can increase its virulence towards *P. guildinii* by growth on HC15, and that exposure to this short chain alkane induces not only genes related to the degradation of epicuticular hydrocarbons but also primes the activity of genes that are needed before or after epicuticle penetration. The virulence of ILB299 was also enhanced but to a lower extent, and in this case, this could be related to the overexpression of genes related to CHC metabolism only. These results suggest that growth on CHC, in a Petri plate, can trigger the expression of genes related to CHC metabolism, adhesion and virulence, and that this early priming can lead to virulence enhancement against insects that are highly resistant to entomopathogens.

## Acknowledgments

This work was funded by INIA Uruguay, Grants SA_24 and SA_47, and Agencia Nacional de Investigación e Innovación (ANII). We thank Sofia Simeto, Gonzalo Martínez and Ximena Cibils for insect rearing.

## Notes

### Competing Interest Statement

The authors have declared no competing interest.

